# Building flexible and robust analysis frameworks for molecular subtyping of cancers

**DOI:** 10.1101/2023.05.01.538907

**Authors:** Christina Bligaard Pedersen, Benito Campos, Neeraja M. Krishnan, Binay Panda, Kristoffer Vitting-Seerup, Maria Rossing, Frederik Otzen Bagger, Lars Rønn Olsen

## Abstract

Molecular subtyping is essential to infer tumor aggressiveness and predict prognosis. In practice, tumor profiling requires in-depth knowledge of bioinformatics tools involved in the processing and analysis of the generated data. Additionally, data incompatibility (e.g., microarray vs. RNA sequencing data) and technical and uncharacterized biological variance between training and test data can pose challenges in classifying individual samples. In this article, we provide a roadmap for implementing bioinformatics frameworks for molecular profiling of human cancers in a clinical diagnostic setting. We describe a framework for integrating several methods for quality control, normalization, batch correction, classification, and reporting and a use case of the framework in breast cancer.

## 1. Introduction

Large-scale multi-omics profiling experiments during the last decade have defined tumor subtypes in glioblastoma, breast cancer, squamous cell lung cancer, and colorectal cancer [1–4]. Morphologically similar tumor samples can differ substantially in underlying genetic aberrations and changes in gene expression. For several cancers, specific tumor subtypes have been linked to different treatment responses and prognoses [5–7], providing an important approach for patient stratification and its application in precision medicine.

In the case of breast cancer, the definition of subtypes has a history extending two decades [8–10] and tumor profiling is already integrated into many modern clinical workflows [11]. More than 100 expression profiles have been proposed [46], however, in practice, the most widely used subtyping methods are the PAM50 [9] and CIT [10] gene signatures. PAM50 is used to classify samples into one of four subtypes: Luminal A, Luminal B, HER2-enriched, and Basal-like. A fifth subtype derived from normal breast tissue was also included in the original work: Normal-like [9]. Classification into these four cancer subtypes was originally based on research microarray and qRT-PCR data, but later an FDA-approved platform was developed [13]. Using a similar approach relying on unsupervised clustering, Guedj et al. (2012) [10] grouped breast cancer samples into six subtypes; lumA, lumB, lumC, mApo, basL, and normL. Additional subtypes, for example, Claudin-low group [14], were discussed as a subtype in the past but later inferred as a phenotype extending across the spectrum of the other groups [15].

Despite their wide use, these subtyping methods are associated with technical challenges in a diagnostic laboratory. For example, most sample sizes in clinical settings are small and may include as few as a single sample from an individual patient. However, the subtyping methods depend on the distribution of data from a group of samples. Additionally, classification results may differ depending on whether a single sample or a batch of samples are used [12]. Additionally, subtype assignment depends on the amount of non-tumor cells in a given sample; and thus, should consider tumor purity. Lastly, many pivotal subtyping methods were derived from microarray data, while most laboratories have moved or are moving towards RNA sequencing for gene expression profiling. The incompatibility of these two data types makes it challenging to leverage established methods while transitioning to more prevalent and unbiased gene expression profiling technologies.

Here, we describe practical bioinformatics solutions to address these challenges with a framework and apply the resulting framework to the PAM50 [9] and CIT [10] classifiers. We provide a generally applicable roadmap for implementing data and bioinformatics tools for robust inference of cancer subtypes in a clinical setting, both from raw and processed RNA sequencing data and on a per-sample basis. Finally, we discuss several other molecular profiling features which, together with subtyping, can be combined into a comprehensive report to support diagnostic and clinical insights.

## 2. Materials and methods

### 2.1. Data

#### TCGA

We used data from the cancer genome atlas (TCGA) for assessing samples in our use case for breast cancer. We downloaded gene expression data from the Xena Browser [16] as transcripts per million (TPM) quantified using RSEM and TOIL, mapped to Gencode GRCh38.p3. For visualization purposes, we selected 20 random samples from each cancer type.

#### GTEx

We downloaded data from the Genotype-Tissue Expression (GTEx) project for assessing whether our use case samples resembled healthy breast tissue. Gene expression data was downloaded from the UCSC Xena project [16] as transcripts per million (TPM) quantified using RSEM and TOIL, mapped to Gencode GRCh38.p3. For the visualizations presented here, we selected 20 random samples from each tissue type.

#### CIT reference data

The CIT microarray (Affymetrix HG-U133 Plus 2.0) training data was downloaded from ArrayExpress (accession: E-MTAB-365) and a subset of data for the 355 core samples defined by the original publication [10]. The .CEL files were read and RMA-normalized using the affy R package [17]. The subtype for each of the samples was available in the data of the citbcmst R package [10] (package no longer maintained).

#### PAM50 reference data

We used the breast invasive carcinoma samples from the TCGA for which a PAM50 subtype has been assigned, to train a classifier for the PAM50 subtyping scheme. The expression of the 50 genes as defined by Parker et al. (2009) [9] were used for classification.

#### Use case data

The use case data set comprises RNA-seq data from the tumors of 57 breast cancer patients sequenced at the Breast Cancer Translational Research Laboratory, Institut Jules Bordet, Brussels, Belgium. It was originally presented by Fumagalli et al. (2014) [18] and derived from EGA (accession number EGAD00001000627). Ten of these samples (named HER2-03, HER2-21, LUMA-18, LUMA-24, LUMA-27, LUMA-29, LUMB-01, LUMB-17, TN-18, and TN-22) were selected to be the use case data set in this study, such that the included samples spanned the four IHC- and grading-based subtypes from the original work (Supplementary Table 1). For CIT classification, RNA sequencing reads were mapped to the Affymetrix HG-U133 Plus 2.0 probe set sequences [19]. For comparison to GTEx, TCGA, and PAM50 classification using TCGA-BRCA as a reference, reads were mapped to Gencode GRCh38.p3.

### 2.2. Methods

#### Quality assessment

We used arrayQualityMetrics v3.54.0 to assess data quality for microarray data [20] and FastQC v0.12.0 [21] for RNA sequencing data.

#### Tumor purity estimates

We estimated tumor purity using a functional class scoring based method, ESTIMATE v1.0.13, which provides enrichment scores for a stromal content gene set and an immune infiltrate gene set in a given sample [22]. These scores are then aggregated into a score that serves as an indicator of tumor purity. As this method works by calculating the enrichment of two gene sets it is important to ensure overlap between the gene symbols in the gene sets and the samples.

#### Data harmonization

Data sets were harmonized using either ComBat [23] (implemented in the SVA package v3.46.0) or simple rank transformation of expression values.

#### Dimensionality reduction and projection

All dimensionality reduction was done using principal component analysis (PCA) with the stats package in R. Initial PCA spaces were constructed on references and additional samples were projected into the reference space by multiplying the rotations from the reference space with the additional sample vectors. For construction of the TCGA and GTEx reference spaces, features were first reduced by subsetting significantly differentially expressed genes between samples from each subtype versus the rest of subtypes collectively, using a Mann-Whitney U test. For the TCGA BRCA reference PCA, the 1092 samples with assigned PAM50 subtypes were subsetted to genes with a log_2_-transformed fold change greater than 1 or smaller than -1. The choice of this filtering setting was based on cross-validation performance. For construction of the CIT PCA reference space, we used the 375 probe sets originally defined by the authors.

#### Definition of subtype-specific gene sets

Subtype-specific gene sets were defined using a Mann-Whitney U test on samples in each subtype versus the rest of the samples. Features were subsetted first based on Bonferroni multiple testing corrected p values lower than 0.05, and then by log_2_-fold changes greater than 1.

#### Classification

Classification of samples was carried out using three different approaches: *k*-Nearest Neighbor (kNN) using the e1071 v1.7-13 package in R, distance-to-centroid, and subtype gene set single sample gene set enrichment using the Singscore package v1.18.0 in R [24]. For the kNN classifier, the best *k* was empirically determined by leave-one-out cross-validation. Then, a winner-takes-all approach was used to assign subtypes. For the distance-to-centroid classifier, centroids were calculated as the mean expression of genes in each class, and the minimum centroid distance subtype was assigned. It has been proposed to shrink the centroid means to offset the effects of outliers on the mean. We observed that defining centroids on the median expression worked just as well. For centroid distance using gene expression, Euclidean distance was used, and for the rank transformed data, we used the Kendall tau distance. For subtype gene set single sample gene set enrichment, we used Singscore. For performance evaluation, we used a leave-one-out approach and calculated precision and recall for each subtype as well as a subtype frequency weighted accuracy.

## 3. Results

### 3.1. Pipeline for molecular subtyping

A robust workflow for subtyping involves several steps (Figure 1). Briefly, the process starts with data preparation where one or more reference data sets (e.g., subtype training samples) and a test data set (new samples to be profiled) are processed. The latter step may also involve some additional steps, to make test data compatible with additional reference sets like the TCGA or GTEx sets. After processing each data set separately, it is necessary to harmonize them to ensure that they can be directly compared, after which the actual subtyping can take place. In this paper, we describe each step of the pipeline and exemplify the analysis using a data set of ten breast cancer samples analyzed with RNA sequencing. However, the workflow is designed to be applicable, in principle, to any other samples, cancers, or data types.

**Figure 1:**
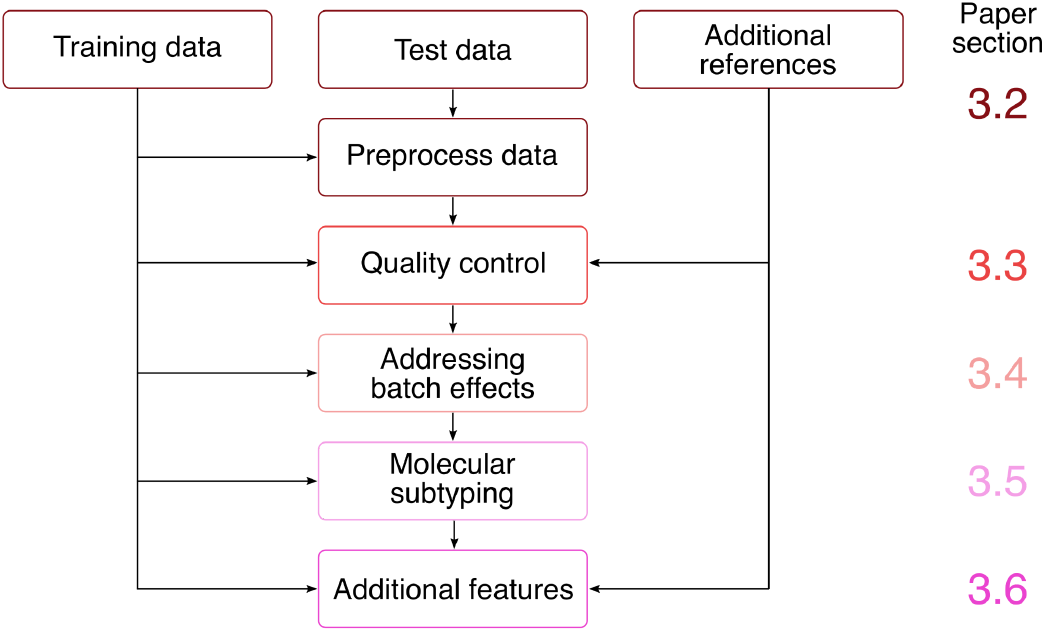
Proposed pipeline for molecular characterization of cancer. The section describing each step of the workflow is indicated on the right.

### 3.2. Preparing data

The first step in building a subtyping framework is to prepare the data set(s) that should serve as a reference for the subtypes. This set will typically be published along with the method describing the subtyping scheme e.g., raw data from the CIT subtyping method published by Guedj et al. (2012) [10]. As a general rule of thumb, the training data should be preprocessed in accordance with the methods used by the authors of the original publication, although this may not affect classification performance in practice.

For samples analyzed with the same Affymetrix platform, extracting the matching probe set intensities is trivial. However, gene expression profiling has largely shifted towards RNA sequencing, as this technique is now both cheaper and more comprehensive. To address this technology incompatibility, we previously devised a method for integrating RNA sequencing data with DNA microarray data [19]. In short, rather than mapping RNA sequencing reads to a reference transcriptome, we propose mapping the reads to the probe set target sequences of the microarray. This also ensures full overlap between the features of the reference set and the test set, which is necessary for harmonization, visualization, and classification. An additional note regarding the use of sequencing data, is that length-normalization is necessary for inter-gene comparisons, on which subtyping is typically highly dependent. One approach is to use transcript per million-normalization.

### 3.3. Quality control

Quality control has two major components: technical and non-technical quality control. Technical quality control strictly deals with the quality of the data from an instrumental point of view. Depending on the platform used, different software packages can be applied to assess the raw data. Data files with poor quality data should be discarded as this can severely affect the results of the downstream analyses. Quality assessment of microarray data can be done using tools such as the Bioconductor package arrayQualityMetrics [20], while RNA sequencing data can be assessed using FastQC [21].

Non-technical quality issues, such as sample impurity or sample swaps can be harder to detect. However, the majority of these non-technical quality parameters can be estimated by comparing sample characteristics to appropriate reference data sets. Tumor purity naturally varies, and may not be an issue for subtyping classification, unless the purity of a sample is outside of the range of the reference set. Tumor purity can be estimated using the software package ESTIMATE [22], which utilizes ssGSEA [25] to calculate enrichment of an immune signature and a stromal signature, which are then aggregated into a tumor purity score. As the method is rank-based it is less sensitive to batch effects. A subtyping analysis of a breast cancer sample with potential low tumor cell content can be done by comparing the purity scores of the samples of interest to the purity scores of all breast cancer samples in a reference set. The ten samples used for exemplifying the framework throughout this paper provide a very broad range of tumor purity (Figure 2).

**Figure 2:**
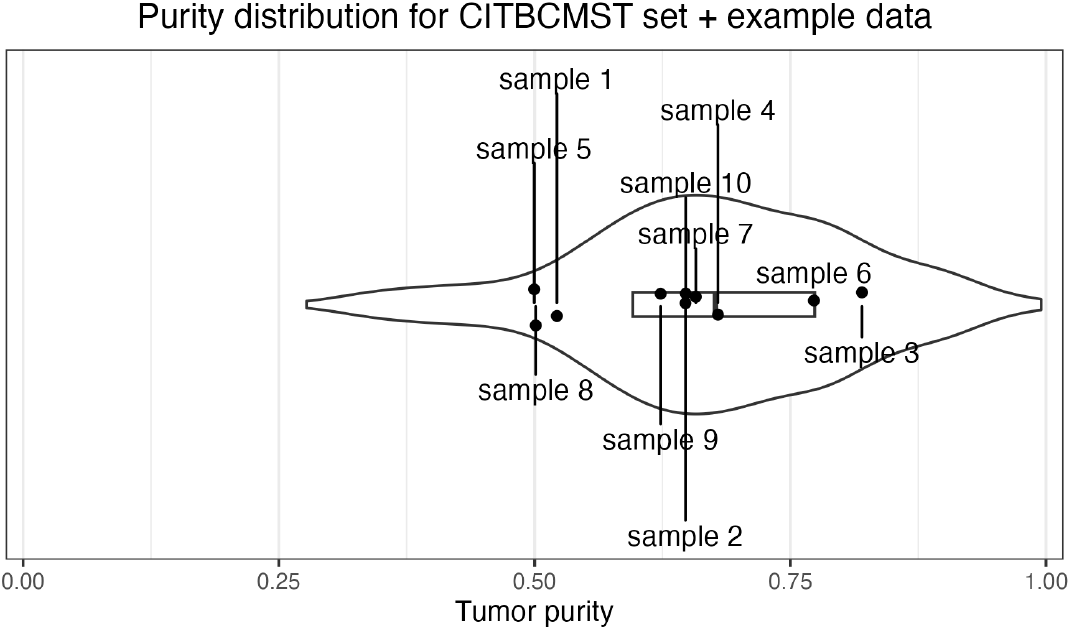
Distribution of purity scores of the CIT reference samples (shown as violin with boxplot) and the ten samples from our test set (shown as labeled points). The x axis represents purity in the range [0; 1]. A lower score indicates lower purity.

Purity reflects the degree to which an expression profile reflects patterns considered to represent normal cells which are also present in tumors including stromal and immune cells, and may need to be explicitly included in tumor expression profiling. For example, two tumor samples which are both characterized by high expression of a specific receptor, but which have different purity, may have very different measured expression values for the given receptor gene based on the tumor sample. Accordingly, if a sample is at the extremes of the purity range, one may consider normalizing expression by purity before reporting.

Generally, for subtyping purposes, it is important to remember that subtypes are defined on bulk samples which include the tumor microenvironment in addition to malignant cells. Accordingly, defined subtypes can actually be correlated with normal cell content, as is the case for both the CIT and PAM50 classifiers (Supplementary Figures 1-2). In some subtyping schemes, the content of non-malignant cells is even the defining feature of specific categories [26].

Sample swaps or major contamination can be difficult to detect, but clues can be provided by comparing sample gene expression to expression characteristics of comprehensive tissue-specific expression profiles such as GTEx [27] or the TCGA [28]. As can be seen in Figure 3, the ten breast cancer test samples appear highly likely to be derived from breast tissue, which correlates with our expectations. Naturally, projection of swapped samples originating from the same tissue into the space of reference sets would not reveal the error. It does, however, indicate whether the analyzed sample actually has an expression profile resembling its true tissue of origin. Furthermore, it may be used to reveal if a sample has been severely impacted by any experimental processing step, e.g., late freezing [29].

**Figure 3:**
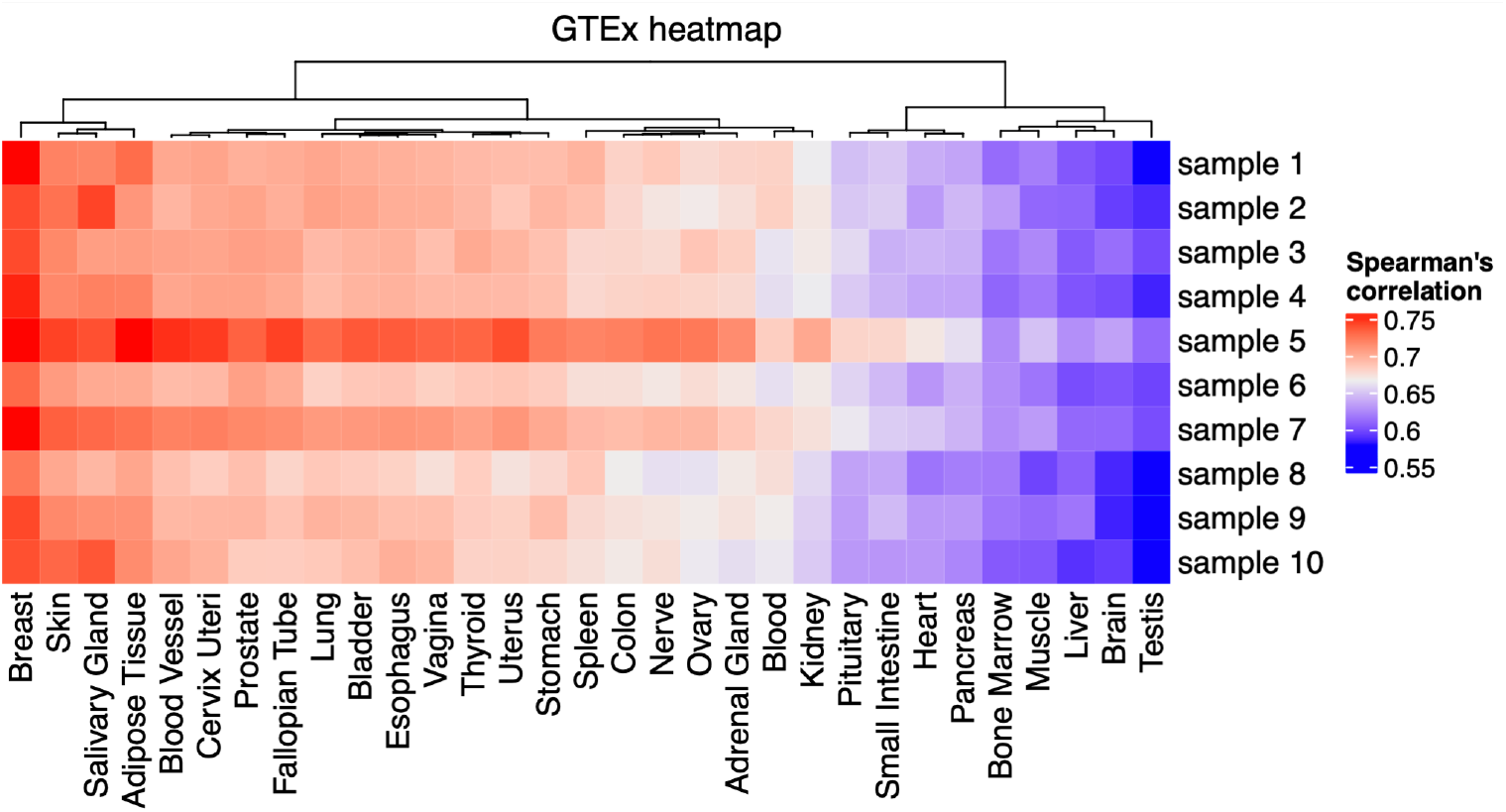
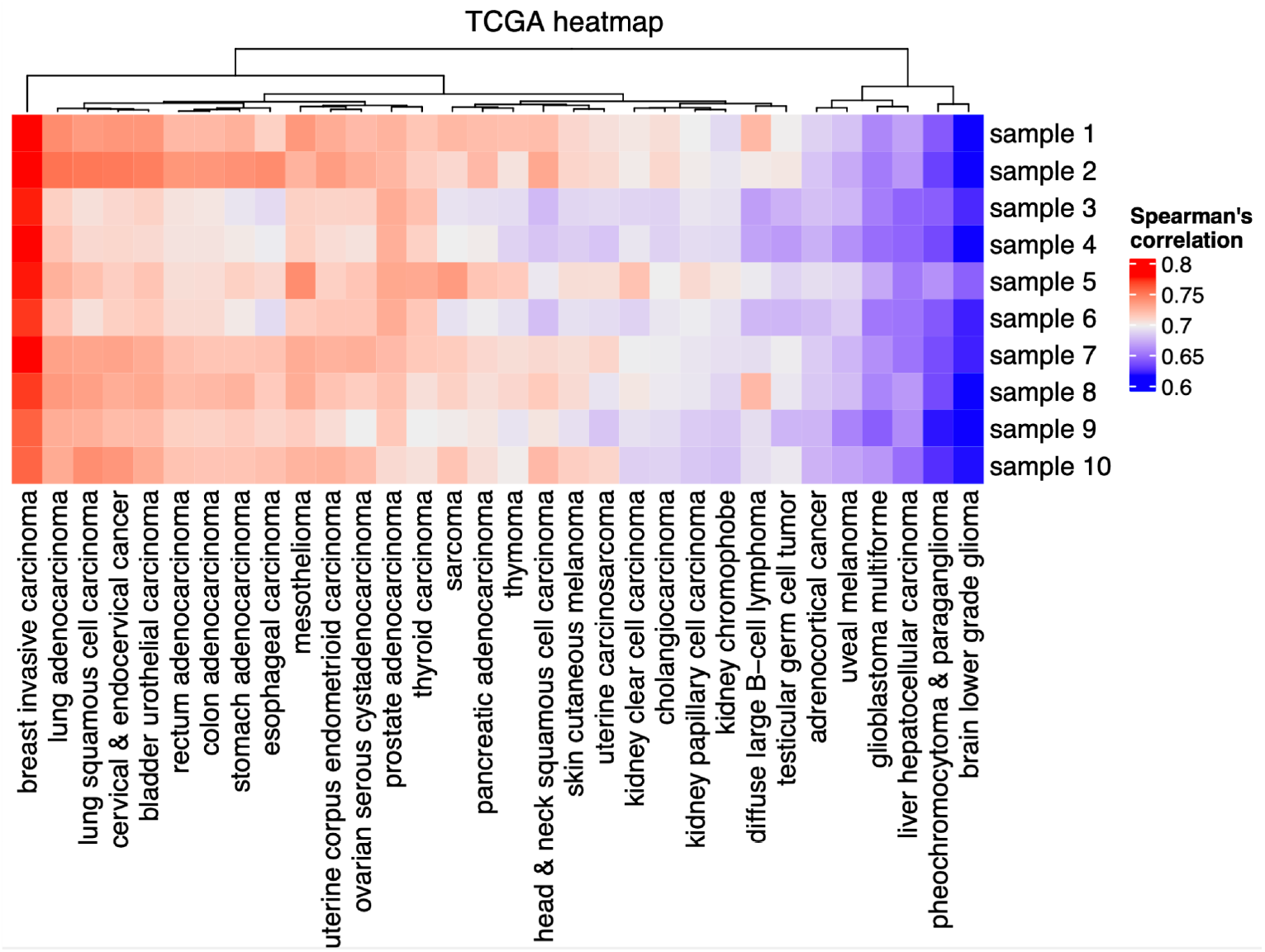
Heatmaps of Spearman’s correlations from samples to tissue mean-based centroids in data sets of comprehensive tissue-specific expression profiles. Columns are clustered and dendrograms are shown. Breast-based classes are highlighted in black squares. A) Heatmap of example samples vs. centroids defined from a collection of healthy tissues from the GTEx project. B) Heatmap of example samples vs. centroids from a collection of cancer tissues from the TCGA project.

### 3.4. Harmonizing data

Before classification, harmonization of expression values from the reference samples is necessary. This is critical both when considering data from the same or very similar platforms, e.g., two microarray experiments. The need for harmonization is even greater when comparing samples across technology platforms. Even after mapping RNA sequencing reads to a reference consisting of probe sets, data from different experiments have completely different distributions, as illustrated in Figure 4A. Harmonization can be carried out using batch correction methods, e.g., ComBat [23] (Figure 4B). While ComBat is designed to work with small sample sizes, at least eight samples must be submitted to adequately model technical variance [23]. If subtyping of fewer samples is needed, expression values can also be rank normalized (Figure 4C).

**Figure 4:**
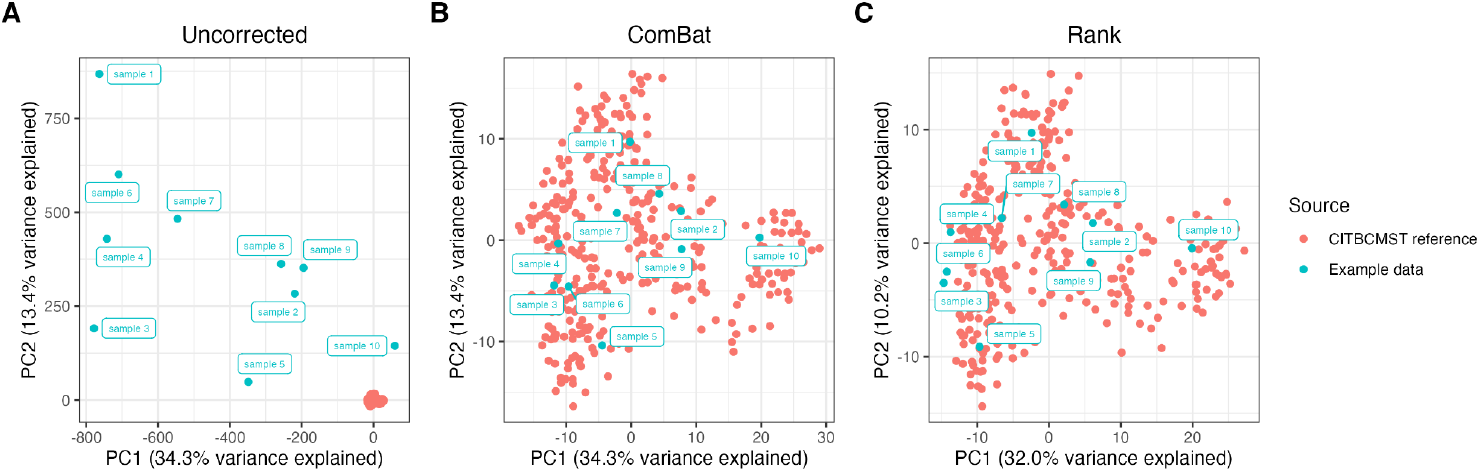
PCA plots of the reference data set (microarray) with the example set (probe-mapped RNA-seq) projected in. Percent variance explained by the PCs refer to the reference data set. A) PCA of samples without batch correction. B) PCA of samples batch corrected with ComBat. C) PCA of samples integrated using rank.

When evaluating batch correction, it is important to consider that the method should provide complete and unbiased integration of the different data sets since even a small shift of the test set can change the distribution of assigned subtypes. ComBat and rank transformation solve the problem of this variation. ComBat explicitly models both technical and biological variance, the latter relying on biological cofactors - i.e., sample condition, or in this case, the hitherto unknown subtype. If batch correcting two large test sets with equal distribution of biological conditions, this may not pose a problem, but if the test data is “biased” towards certain conditions, e.g., includes only one subtype, ComBat is not the optimal solution as illustrated in Figure 5A. Rather, correction methods for which the sample-wise transformation is independent of the other samples in the test set is desirable [12,30], and as can be seen in Figure 5B, rank transformed data is readily comparable to the reference. In situations where the reference data are not provided as raw data, or only include a subset of features, it may be useful to use a signature enrichment approach such as biological process transformation.

**Figure 5:**
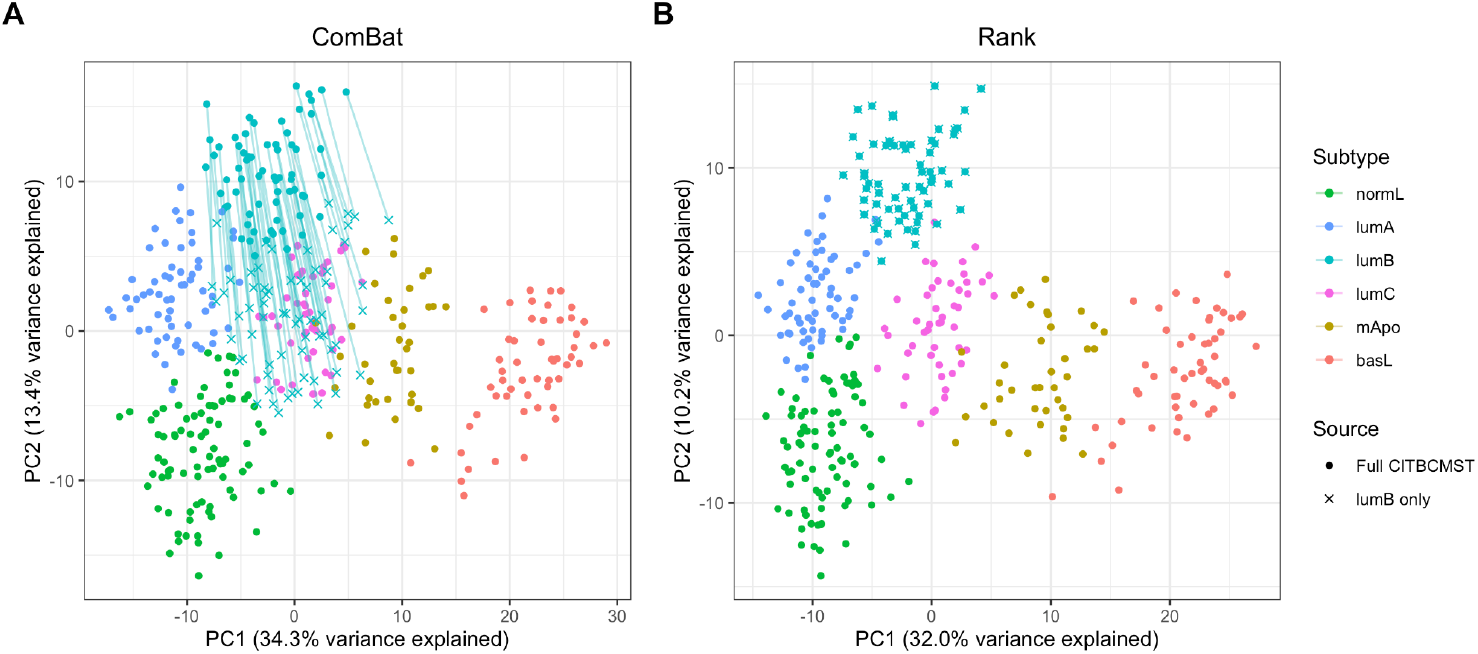
Examples of harmonization of data sets with unbalanced biological conditions. Percent variance explained by the PCs refer to the full CIT set. A) Batch correction of the entire CIT training set against the lumB samples only using ComBat with the full set as reference. Without biological conditions as cofactors, ComBat under-estimates the biological variance causing the lumB samples to become centered in the PC space. Similar plots for the remaining subtypes are shown in Supplementary Figure 3. B) A similar example using rank transformation.

### 3.5. Subtype classification

Once the reference and test data are harmonized, subtypes can be assigned to the test samples. Some subtyping studies provide a software package for classifying new samples, e.g., the CIT classifier for breast cancer [10], while others do not. In the latter cases, an appropriate classifier must be trained and applied. There are no one-size-fits-all approaches for this, and hence, a classifier may be selected based on its cross-validation performance. To demonstrate how this is carried out, we have applied the CIT reference data and leave-one-out cross-validation to test two classification algorithms: a *k*-nearest neighbor classifier and a distance-to-centroid classifier. Advantages of these algorithms are 1) they are quite simple and thus the results are easily interpretable, and 2) subtype assignment probability can be inferred (fuzzy classification), rather than the methods providing a “winner-takes-all” classification. Distance-to-centroid additionally allows for easy outlier detection. Another strategy for classification is to define subtype specific gene expression signatures using ssGSEA based on the training set, and calculate the enrichment score of each signature for each additional sample. An advantage of this approach is that it is less sensitive to missing features resulting from data sparsity (as observed in single cell transcriptomics). As seen in Table 1, all three classifiers perform reasonably well, with a slight performance advantage for the distance-to-centroid classifier.

**Table 1:** Precision, recall, and weighted accuracy from leave-one-out cross-validation of k-nearest neighbor, nearest centroid, and subtype signature enrichment for the CIT reference data. Highest precision per subtype is highlighted in green, highest recall is highlighted in blue, highest weighted accuracy is highlighted in purple.

|  | *k*-nearest neighbor |  | Distance-to-centroid |  | Subtype signature ssGSEA |  |
| --- | --- | --- | --- | --- | --- | --- |
| Subtype | Precision | Recall | Precision | Recall | Precision | Recall |
| **normL** | 0.966 | 0.989 | 1 | 1 | 0.827 | 0.920 |
| **lumA** | 0.909 | 0.984 | 1 | 0.984 | 0.875 | 0.918 |
| **lumB** | 0.955 | 0.955 | 0.957 | 1 | 0.844 | 0.985 |
| **lumC** | 1 | 0.854 | 1 | 0.958 | 0.722 | 0.542 |
| **mApo** | 1 | 1 | 1 | 1 | 1 | 0.795 |
| **basL** | 1 | 1 | 1 | 1 | 1 | 0.925 |
| Weighted accuracy | 0.963 |  | 0.990 |  | 0.847 |  |

The performance of the leave-one-out classification of the reference data set reveals that if reference data and test data is appropriately harmonized, even simple classifiers can perform quite well. Of course, some methods will be preferable to others as illustrated above, but this can easily be established using cross-validation. This conclusion is not only applicable for the specific case of the CIT tool for breast cancer but the approach may be used in much wider contexts across data sets, subtyping schemes, and cancer types.

As mentioned, many classification methods, such as those used here, have the added advantage of fuzzy classification. This means that samples with almost equal distances to two or more subtypes may be considered as mixed cases [10]. In clinical diagnostic settings, such information may be of interest, but if more simple classification is preferred, the single closest centroid can also be reported as winner-takes-all. An example of fuzzy classification is shown in Figure 6, where the ten test samples are classified according to the CIT scheme, using rank transformation followed by a distance-to-centroid classifier. While some samples clearly resemble a single subtype (e.g., sample 10), the label of others is more unclear (e.g., sample 6).

**Figure 6:**
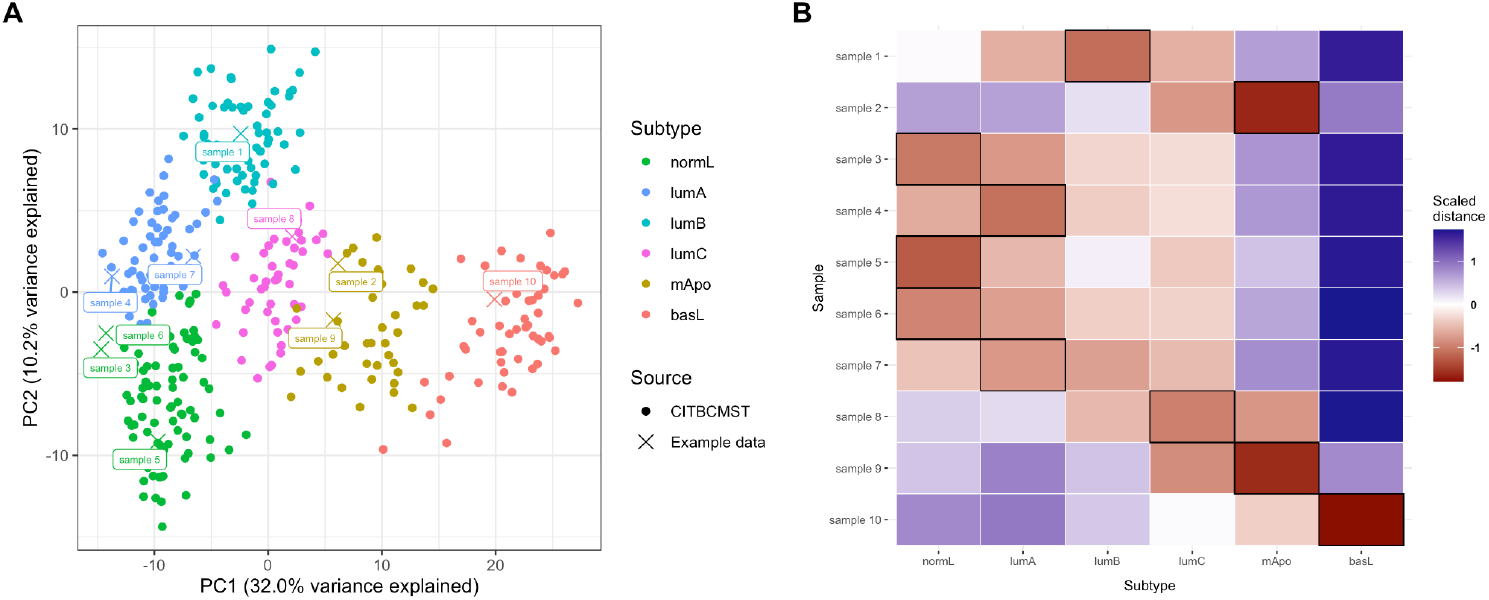
Subtyping of ten test samples using rank transformation and a distance-to-centroid classifier. A) Projection of the samples from the use case data into CIT principal component space. Percent variance explained by the PCs refer to the CIT set. B) Distance-to-centroid matrix, scaled per sample. Black squares indicate the closest centroid for each sample.

A similar approach to classification can be taken for PAM50. The primary difference between CIT and PAM50, is that the former study provided a training set, a feature set, and a classifier, while the latter provided a 50-gene signature. Using the TCGA BRCA data with the ranks of the 50 PAM50 genes, a nearest centroid classifier yields a weighted accuracy of 0.894 (See Supplementary Table 2 for full performance metrics). As shown in Figure 7, a majority of our included samples are labeled as LumA. While this fits nicely with the IHC-based ER-positive status of all of these six samples, it is somewhat conflicting that sample 1, which is also HER2-positive, is not classified as Her2, but this sample was similarly classified as luminal with CIT. In the case of sample 7, it is also quite close to the LumB subtype (Figure 7b), so this could be considered as a mixed assignment. Sample 2, sample 8, and sample 10 behave as expected from their receptor status, but the assignment of sample 9 to the Normal-like subtype may seem unexpected. Of note, the PAM50 definition of Normal-like is actually based on non-cancerous breast tissue samples but the assignment of receptor-negative samples to the Normal-like PAM50 subtype is not unusual [31].

**Figure 7:**
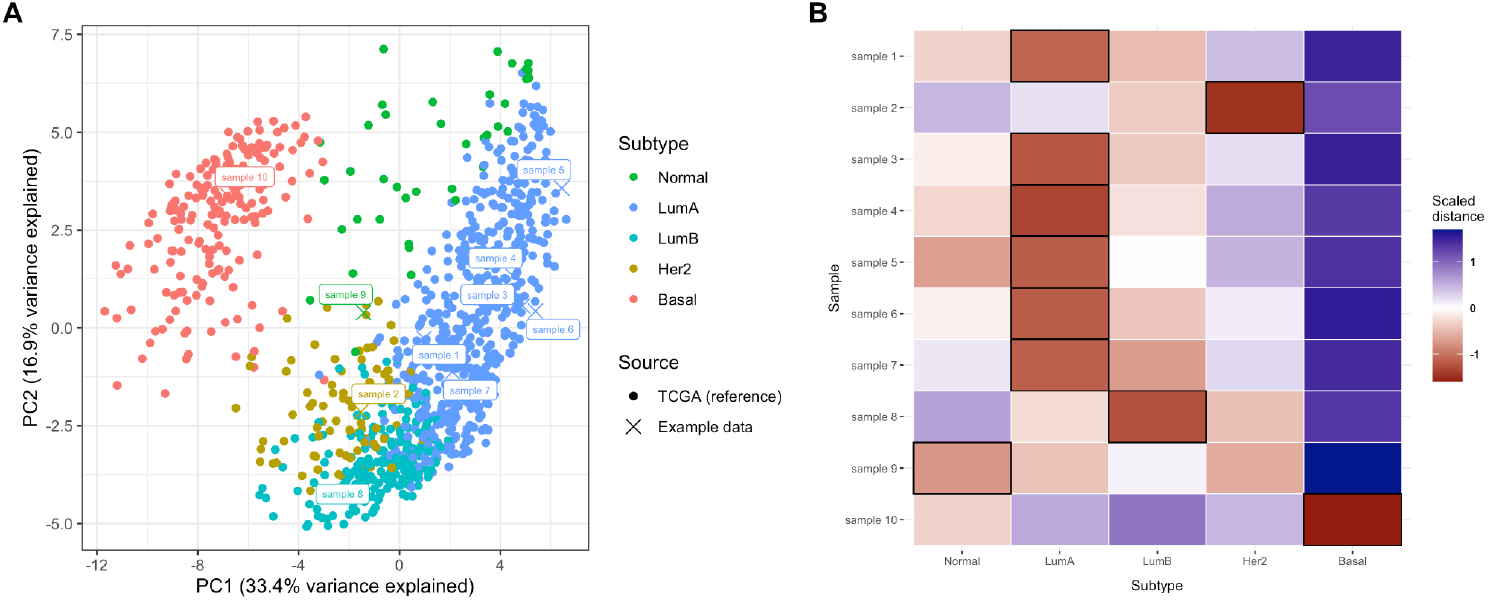
Subtyping of ten test samples using rank transformation and a distance-to-centroid classifier. A) Projection of the samples from the use case data into PAM50 principal component space. Percent variance explained by the PCs refer to the TCGA reference set. B) Distance-to-centroid matrix, scaled per sample. Black squares indicate the closest centroid for each sample.

### 3.6. Additional features

Besides the assignment of subtypes based on gene expression analysis, additional tumor features may be of interest when optimizing treatment regimens. In the case of breast cancer, this includes expression levels of specific receptor genes, such as the estrogen receptor (ER), progesterone receptor (PR), and the HER2-receptor [32]. While receptor status is mainly based on immunohistochemistry or ERBB2 FISH probes, multiple studies have shown that measurements at the mRNA-level have reasonably strong correlations with those from IHC [10,32–34]. Of note, a sample’s expression level of any tumor-associated gene may be impacted by the sample purity, as also described in brief in Section 3.3.

Generally speaking, any feature may be visualized in relation to any reference (e.g., subtyping reference, TCGA, in-house samples, etc.), as seen in Figure 8. In addition to genes, enrichment of relevant gene signatures or pathways related to prognostics can also be visualized. More widely across cancer types, this can include detection of specific mutations [35,36], or methylation analyses [37]. Such analyses add to the complexity and costs of molecular profiling, but can also provide valuable information.

**Figure 8:**
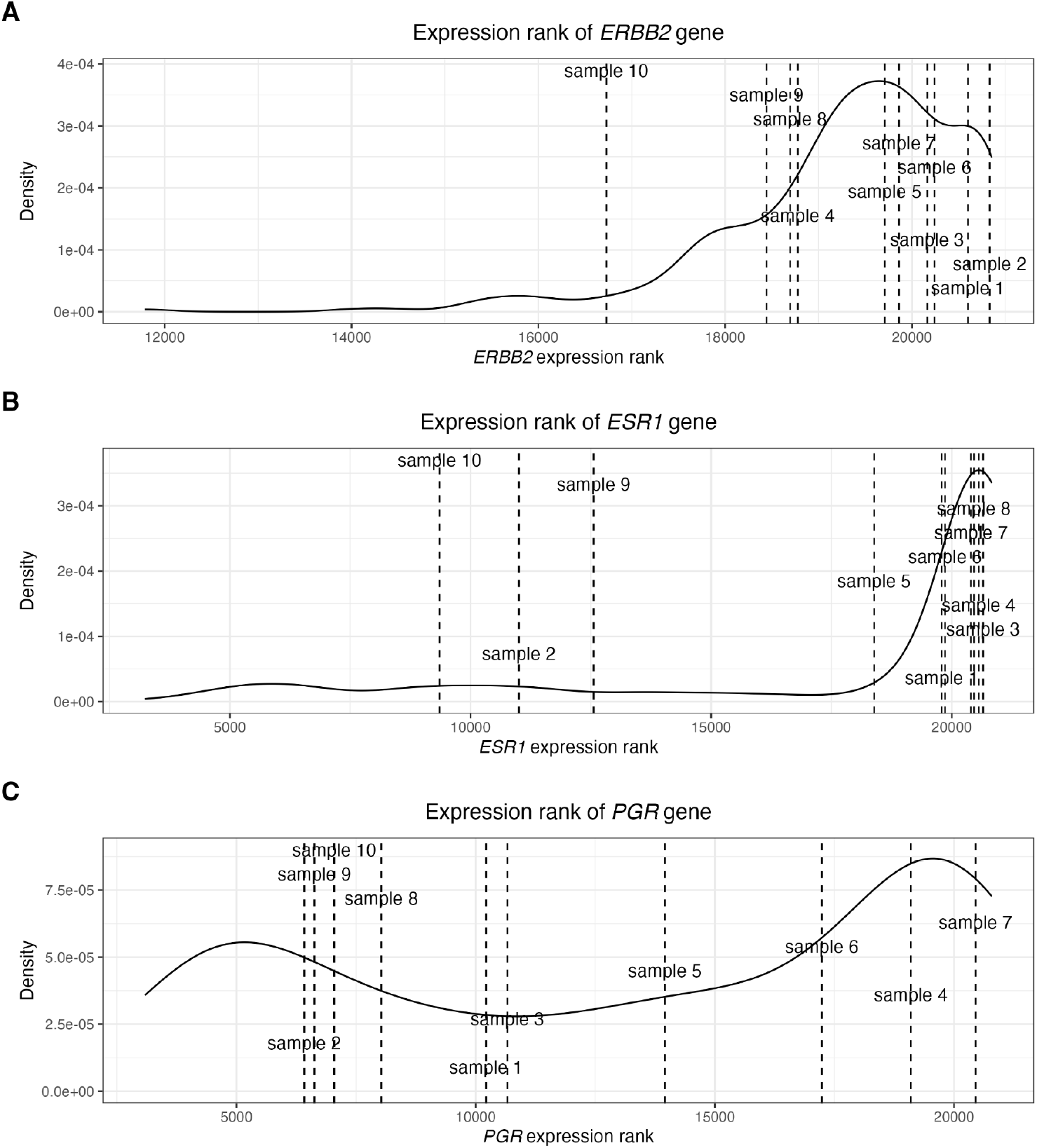
Visualizing the expression ranks of specific genes of interest in relation to the expression rank distributions in the reference data (CIT). A) Expression of *ERBB2* (HER2). B) Expression of *ESR1* (Estrogen receptor). C) Expression of *PGR* (Progesterone receptor).

## 4. Discussion

Molecular characterization of tumors is an important tool for precision medicine. In this paper, we describe a bioinformatics framework based on years of practical experience. Most characteristics rely on comparisons to reference samples. For tumor purity, the sample can be compared to the previously collected and analyzed samples to establish whether the sample is within a reasonable range. For quality control, comparisons to different cancer samples in the TCGA data or the healthy tissues in GTEx may reveal more serious sample issues, if samples do not resemble the expected tissues.

For subtyping, samples are directly compared to the reference data sets. For these comparisons to yield sensible results, mapping, normalization, and harmonization must be considered, as transcriptomic measurements are sensitive to batch effects and technology incompatibility (e.g., comparing microarray data and RNA sequencing data). In the case of CIT (and many other subtyping methods), gene expression profiles were measured using the Affymetrix HG-U133 Plus 2.0 DNA microarray. In the original study, the subtyping itself was not based on summarized gene expression values, but a selection of probe set intensities. This means that while gene-level classification is possible using the classifier offered by the authors of CIT, additional samples should optimally be classified based on intensities of the same probe sets. Consequently, for comparing RNA sequencing data to microarray, mapping to probe set sequences greatly increases compatibility.

The batch correction tool of choice in the vast amount of transcriptomics applications is ComBat. However, ComBat is dependent on biological cofactors to accurately model biological variance. Therefore, using ComBat carries the risk of removing biological variance without knowing the subtypes in advance. Instead, it is recommended [30,38,39] to use the expression ranks of genes. One caveat is that gene expression ranks may be sensitive to noise - particular in the low end of the expression spectrum, where even a small change can induce swaps in the ranks. One option is to apply the robust rank aggregation [40], although when working with small, curated gene sets such as PAM50, the effect is negligible.

Once data is harmonized, dimensionality reduction and classification can be performed on the ranks. For distance-based classifiers, a distance metric suitable for ranks should be selected. Some studies have presented advanced machine learning classifiers for breast cancer samples [41–43]. In the present paper, we intentionally focused on three very simple methods, and show that with proper data harmonization, even simple models will work. Simple models come with the added bonus of interpretability and are widely useful, especially in resource constrained settings. Our classifiers were based on the core gene sets defined by the original studies. Others define their own gene sets to optimize cross-validation performance [12,42]. While neither approach is methodologically wrong, it does open the debate of whether the subtype calls or the genetic features on which they are based are of greater importance.

One of the classification methods we chose to apply, stands out from the rest: the subtype-specific single sample gene set enrichment analysis. Though it was the poorest performing of the three tested methods, we chose to highlight this method as it is more robust to data sparsity [44], which may prove important in single cell diagnostics [45]. One important thing to consider in this context is that signatures derived from bulk transcriptomics include signals from the entire microenvironment, and as we show here, tumor purity not only impacted the definitions of the subtypes, but also the subsequent molecular characterizations. This means that single tumor cells may not readily fit into current subtyping schemes.

For diagnostic applications, stability is essential. This means that the stability of the software packages used, R versioning, etc. must be taken into consideration. This is essential both from an operational point of view, as well as from an analytical point. This may be addressed by using a docker environment.

## 5. Conclusion

In this paper, we have presented a framework which enables robust molecular characterization of clinical cancer samples. Particularly, the work emphasizes that harmonization of the new data to be classified relative to the reference is essential for deriving a molecular subtype. The method for harmonization is important. Methods relying on intra-sample relative expression values are particularly suited for classification of single samples. We showed that once data is properly harmonized, even simple classification models can yield high accuracy.

## Author contributions

Conceptualization, LRO, BC and CBP; Methodology, LRO, CBP, KVS, FOB; Software, LRO and CBP; Formal Analysis, LRO and CBP; Investigation, CBP, LRO, MCR, FOB; Resources, LRO; Data Curation, LRO, CBP, BP, NMK BC, MCR, FOB; Writing – Original Draft Preparation, CBP, LRO, MCR; Writing – Review & Editing, CBP, BC, NMK, BP, KVS, MCR, FOB; Visualization, CBP, LRO; Supervision, LRO, MCR; Project Administration, LRO; Funding Acquisition, LRO

## Data availability statement

All data is available via public repositories. TCGA BRCA and GTEx data was downloaded from Xenabrowser.net (dataset ID: TcgaTargetGtex_rsem_gene_tpm). The CIT microarray (Affymetrix HG-U133 Plus 2.0) training data was downloaded from ArrayExpress (accession number: E-MTAB-365). The use case data set is available in EGA (accession number: EGAD00001000627).

