## Supplementary figures and tables for "Building flexible and robust analysis frameworks for molecular subtyping of cancers"

### Supplementary materials

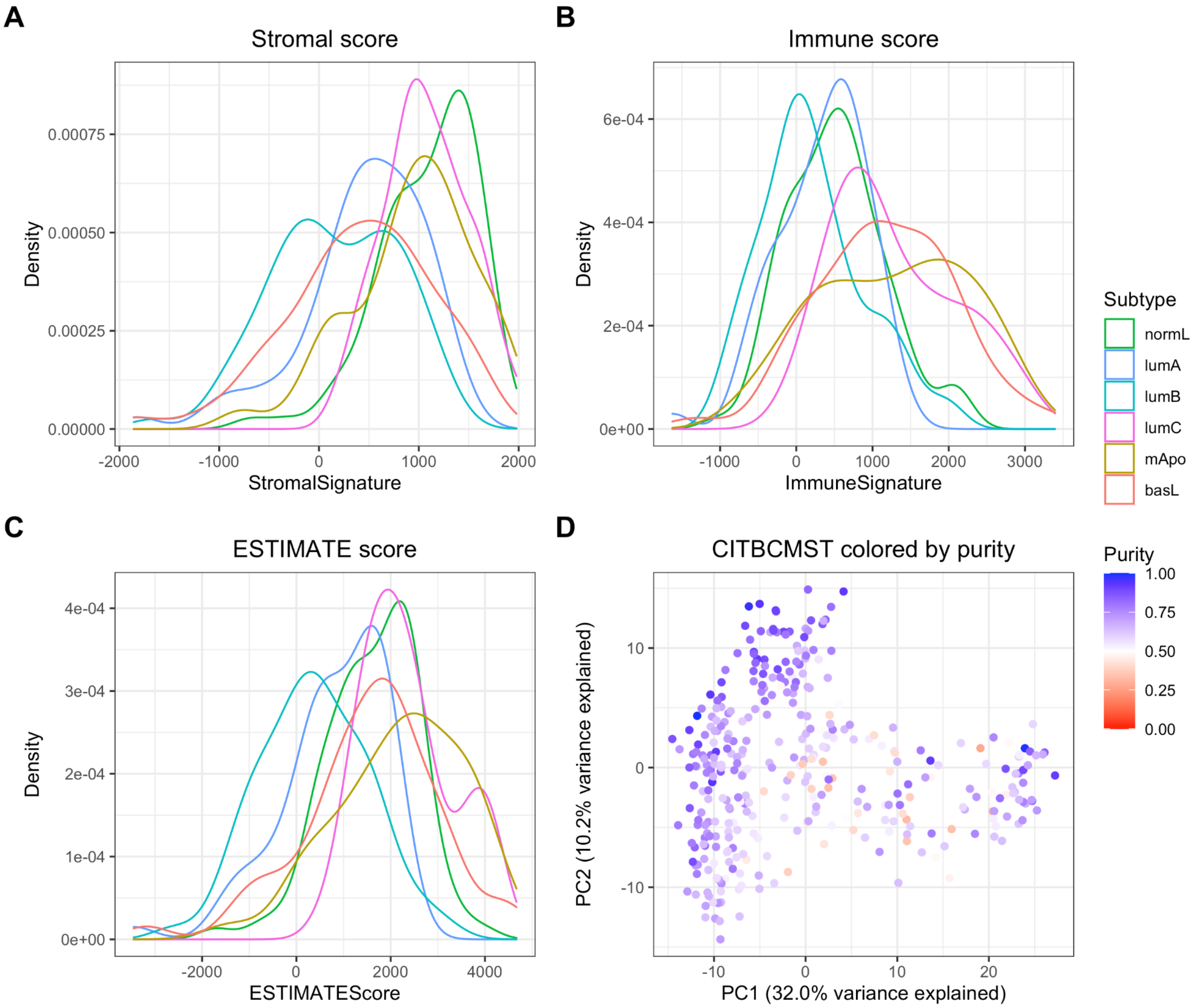


Supplementary Figure 1: Purity estimates for the CITBCMST training data. A) Stromal score distribution for each of the six subtypes. B) Immune score distribution for each of the six subtypes. C) ESTIMATE score distribution for each of the six subtypes. D) PCA plot of ranked CITBCMST training data colored by estimated tumor purity.

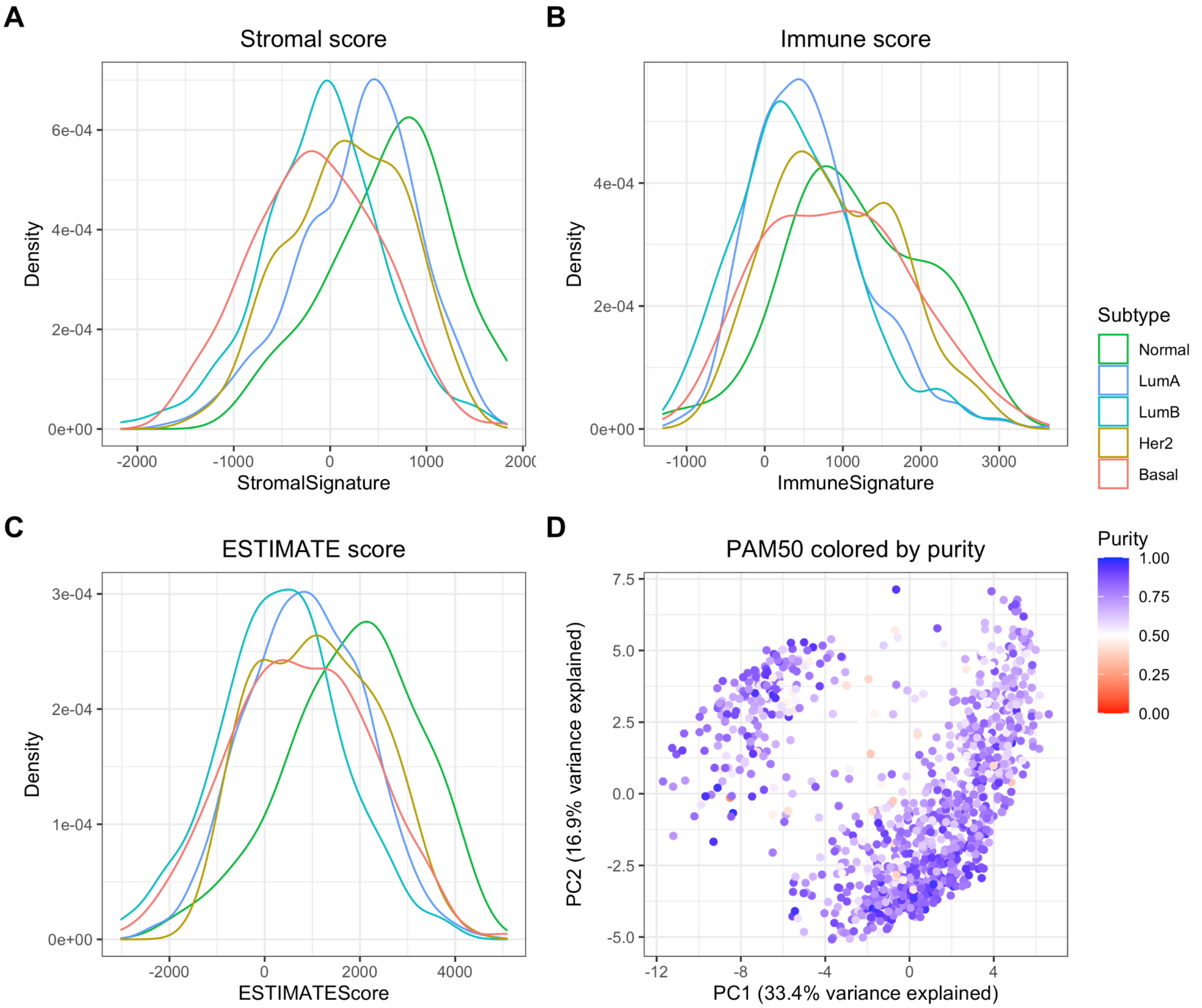


Supplementary Figure 2: Purity estimates for the TCGA data. A) Stromal score distribution for each of the six subtypes. B) Immune score distribution for each of the six subtypes. C) ESTIMATE score distribution for each of the six subtypes. D) PCA plot of ranked TCGA data colored by estimated tumor purity.

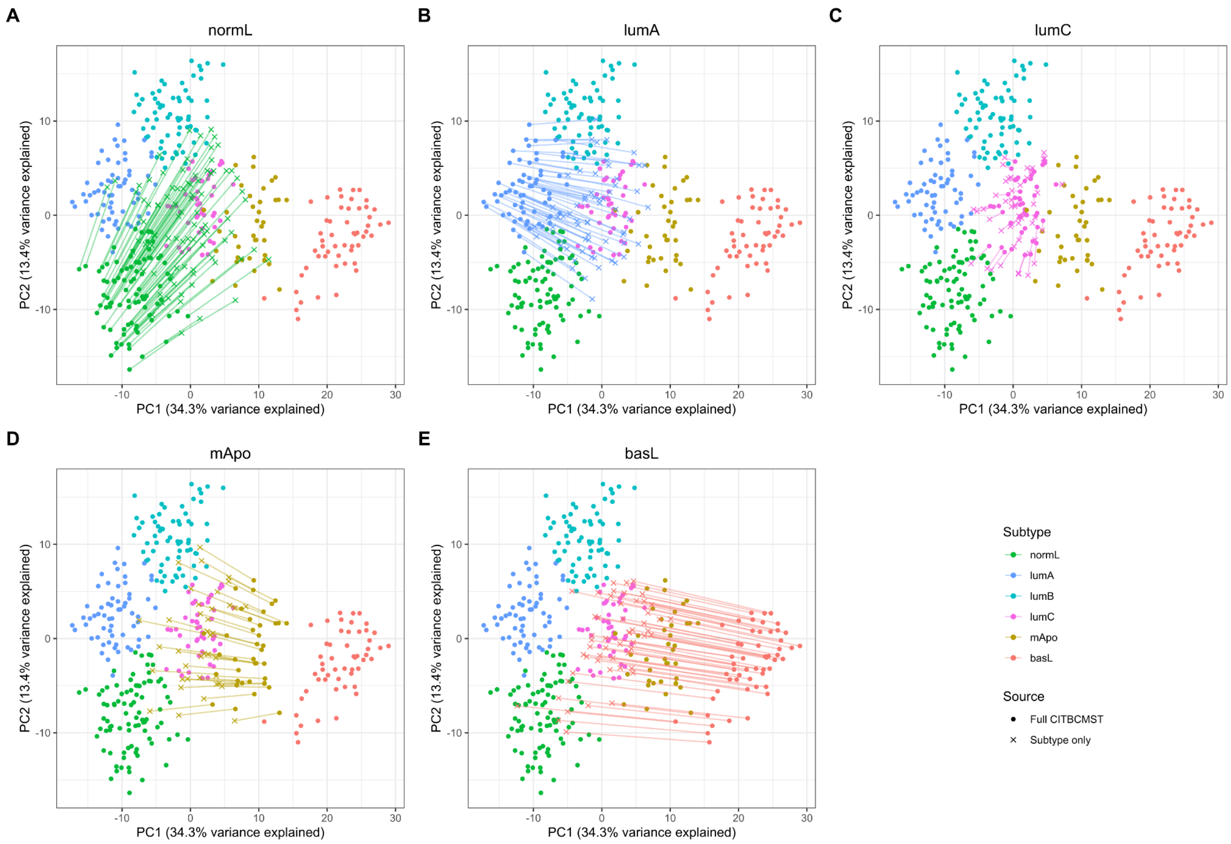


Supplementary Figure 3: ComBat on five subtypes individually and how it affects the location in PC space. Batch correction of the entire CITBCMST training set against the samples from each subtype alone, using ComBat with the full set as reference. Results for lumB are shown in Figure 5. A) normL, B) lumA, C) lumC, D) mApo, E) basL.

Supplementary Table 1: Example patient characteristics.

| Sample | Original name | ER status | PR status | HER2 status | Ki67 % | Grade |
|---|---|---|---|---|---|---|
| Sample 1 | HER2-03 | Positive | Positive | Positive | 20 | 3 |
| Sample 2 | HER2-21 | Negative | Negative | Positive | 70 | 3 |
| Sample 3 | LUMA-18 | Positive | Positive | Negative | 5 | 1 |
| Sample 4 | LUMA-24 | Positive | Positive | Negative | 10 | 1 |
| Sample 5 | LUMA-27 | Positive | Positive | Negative | 5 | 1 |
| Sample 6 | LUMA-29 | Positive | Positive | Negative | 10 | 1 |
| Sample 7 | LUMB-01 | Positive | Positive | Negative | 30 | 3 |
| Sample 8 | LUMB-17 | Positive | Negative | Negative | 10 | 3 |
| Sample 9 | TN-18 | Negative | Negative | Negative | 30 | 2 |
| Sample 10 | TN-22 | Negative | Negative | Negative | 50 | 3 |

Supplementary Table 2: Precision, recall, and weighted accuracy from leave-one-out cross-validation of kNN, nearest centroid, and subtype signature functional class scoring for the PAM50-classification of the TCGA reference data. Highest precision per subtype is highlighted in green, highest recall is highlighted in blue, highest weighted accuracy is highlighted in purple.

| | ***k*-nearest neighbor** | | **Distance-to-centroid** | | **Subtype signature ssGSEA** | |
|---|---|---|---|---|---|---|
| Subtype | Precision | Recall | Precision | Recall | Precision | Recall |
| **Normal** | 0.814 | 0.564 | 0.368 | 0.897 | 0.472 | 0.897 |
| **LumA** | 0.914 | 0.950 | 0.992 | 0.669 | 0.842 | 0.968 |
| **LumB** | 0.853 | 0.829 | 0.624 | 0.949 | 0.923 | 0.552 |
| **Her2** | 0.871 | 0.839 | 0.806 | 0.975 | 0.890 | 0.901 |
| **Basal** | 0.984 | 0.978 | 0.989 | 0.978 | 1 | 0.826 |
| Weighted accuracy | 0.832 | | 0.894 | | 0.829 | |
